# Gene Transfer to the Outflow Tract

**DOI:** 10.1101/044396

**Authors:** Yalong Dang, Ralitsa Loewen, Hardik A. Parikh, Pritha Roy, Nils A. Loewen

**Author notes:** corresponding author: Nils A. Loewen, MD, PhD, 203 Lothrop St, Suite 819, Pittsburgh, PA 15213.

## Abstract

Elevated intraocular pressure is the primary cause of open angle glaucoma. Outflow resistance exists within the trabecular meshwork but also at the level of Schlemm’s canal and further downstream within the outflow system. Viral vectors allow to take advantage of naturally evolved, highly efficient mechanisms of gene transfer, a process that is termed transduction. They can be produced at biosafety level 2 in the lab using protocols that have evolved considerably over the last 15 to 20 years. Applied by an intracameral bolus, vectors follow conventional as well as uveoscleral outflow pathways. They may affect other structures in the anterior chamber depending on their transduction kinetics which can vary among species when using the same vector. Not all vectors can express long-term, a desirable feature to address the chronicity of glaucoma. Vectors that integrate into the genome of the target cell can achieve transgene function for the life of the transduced cell but are mutagenic by definition. The most prominent long-term expressing vector systems are based on lentiviruses that are derived from HIV, FIV, or EIAV. Safety considerations make non-primate lentiviral vector systems easier to work with as they are not derived from human pathogens. Non-integrating vectors are subject to degradation and attritional dilution during cell division. Lentiviral vectors have to integrate in order to express while adeno-associated viral vectors (AAV) often persist as intracellular concatemers but may also integrate. Adeno-and herpes viral vectors do not integrate and earlier generation systems might be relatively immunogenic. Nonviral methods of gene transfer are termed transfection with few restrictions of transgene size and type but often a much less efficient gene transfer that is also short-lived. Traditional gene transfer delivers exons while some vectors (lentiviral, herpes and adenoviral) allow transfer of entire genes that include introns. Recent insights have highlighted the role of non-coding RNA, most prominently, siRNA, miRNA and lncRNA. SiRNA is highly specific, miRNA is less specific, while lncRNA uses highly complex mechanisms that involve secondary structures and intergenic, intronic, overlapping, antisense, and bidirectional location. Several promising preclinical studies have targeted the RhoA or the prostaglandin pathway or modified the extracellular matrix. TGF-β and glaucoma myocilin mutants have been transduced to elevate the intraocular pressure in glaucoma models. Cell based therapies have started to show first promise. Past approaches have focused on the trabecular meshwork and the inner wall of Schlemm’s canal while new strategies are concerned with modification of outflow tract elements that are downstream of the trabecular meshwork.

## 1. Introduction

Open angle glaucoma (OAG), the most common form of glaucoma, is characterized by a decreased outflow facility and elevated intraocular pressure (IOP) (Stamer and Acott, 2012). Progressive loss of retinal ganglion cells is the clinical consequence in all glaucomas and the leading cause of irreversible vision loss worldwide. IOP remains the major modifiable cause and risk factor (Caprioli and Varma, 2011). Compared to traditional interventions that consist of frequently used drops (Friedman et al., 2007), laser (Kaplowitz et al., 2015; Lin, 2008) or filtering surgery (Bussel et al., 2014), gene and cell based therapies have potential to address OAG in a more specific yet long lasting fashion that may also be less traumatic. Gene therapy changes expression in the target tissue by introducing exogenous nucleic acids such as DNA, mRNA, small interfering RNA (siRNA), microRNA (miRNA), or antisense oligonucleotides. Both coding and, more recently, non-coding nucleic acid (Husso et al., 2014) have been used. Cell based therapies often involve transplantation of cells that have been engineered with gene therapy tools. The eye makes for a good target because it is a relatively confined anatomic space that can be directly observed.

Gene transfer using viral vectors has shown promise for many eye diseases but compared to retinal disorders, there is a noticeable paucity of gene therapy trials for glaucoma. Of 2210 clinical gene therapy trials, only 31 (1.4%) were concerned with ocular diseases (“Gene Therapy Clinical Trials Worldwide,” n.d.). Twenty-two of those are based in the United States, two in France (<u>FR-0059, FR-0060</u>), two in Australia (<u>AU-0025, AU-0029</u>), one in the United Kingdom (<u>UK0192</u>), one in China (<u>CN-0025</u>), and one in Israel (<u>IL-0008</u>). There is presently only a single clinical glaucoma gene therapy trial, <u>US-0589</u>, which involves a subconjunctivally injected adenoviral vector prior to trabeculectomy. In this phase 1 trial, p21, a potent cyclin-dependent kinase inhibitor, is expressed to reduce scar formation.

This review will provide an overview of gene transfer to the primary site of glaucoma pathogenesis, the outflow tract, and discuss what has been accomplished and which approaches have not yet been attempted.

## 2. Established methods of ocular gene therapy

In gene therapy, genes can be added, altered, or knocked down by viral vectors, a process termed transduction, or by non-viral methods, termed transfection. Delivery may be directly in vivo or in vitro to cultured cells that are then transplanted. Although most cells in the eye are nondividing, not all vectors can transduce them. Predominantly used viral vectors are based on adenovirus (AdV), adeno-associated virus (AAV), herpes simplex virus (HSV), type C-retrovirus, and lentivirus (Fig. 1). The most important distinguishing features are 1) transgene persistence, 2) packaging capacity, 3) cell type preference, 4) immunogenicity, and 5) safety. Transgene persistence depends on whether or not genomic integration is achieved since episomes or concatemers that contain the transgene are subject to degradation or dilutional attrition by cell division. Transgene integration is by definition mutagenic and may result in unpredictable consequences (disruption of host genes, expression of downstream genes). The packaging capacity is sufficient in most vectors to deliver exons of no more than 2 kilo-basepairs (kb). More complex genes or entire introns, in contrast, can only be packaged by some vectors as reviewed below. Cell type preference depends on transduction kinetics (affected by contact time, transducing units per volume, media) and receptor tropism. Immunogenicity is influenced by viral proteins, prior exposure, transgene species, and cell lines and media used during production (Bessis et al., 2004). The biosafety level is especially concerned with tropism of the natural host (human versus nonhuman), creation of aerosol, and potential for a replication-competent vector that might start to propagate.

**Figure 1.**
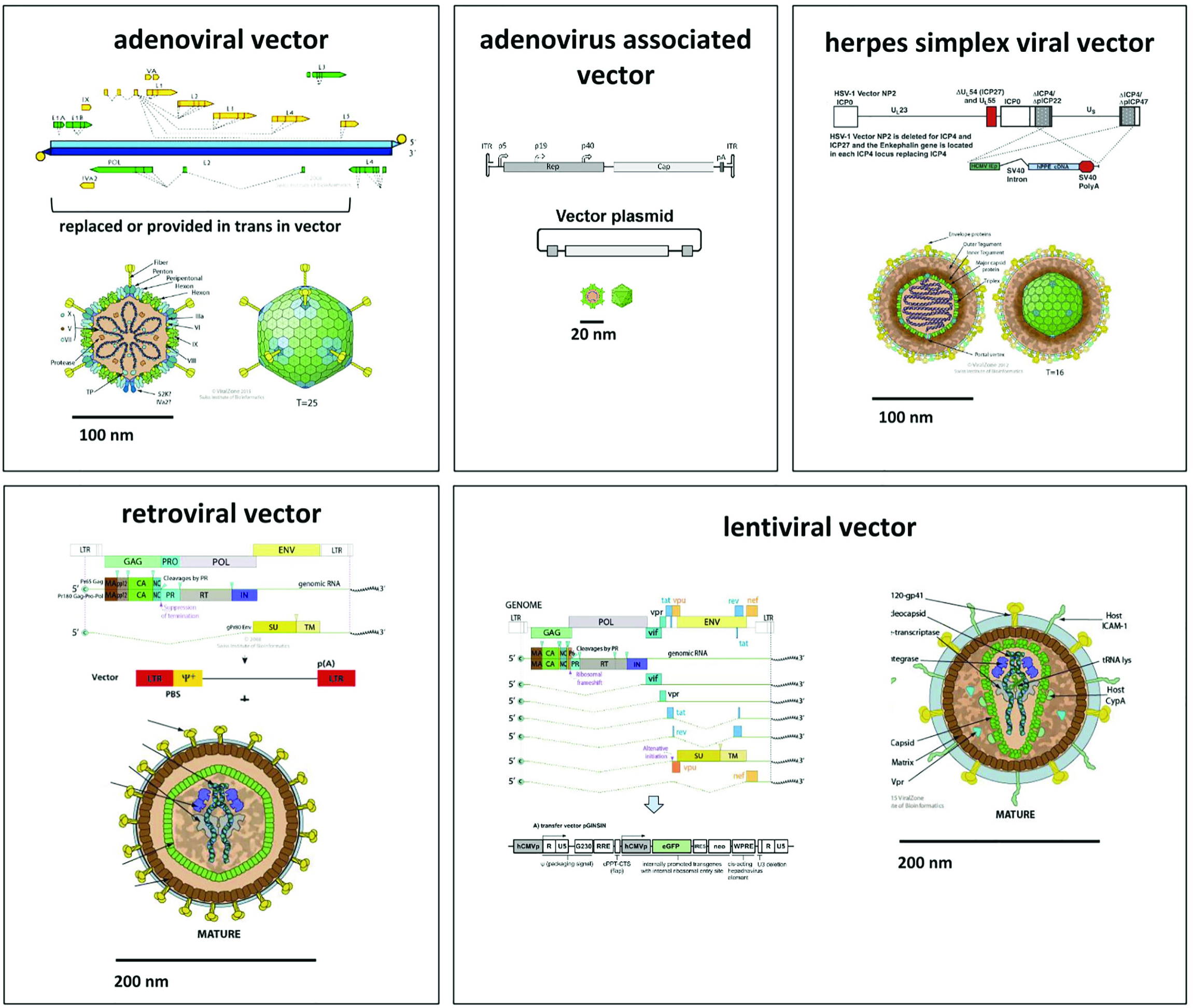
Fig. 1. Genome and structure models of vectors commonly used in ocular gene transfer. Adenoviral vectors are non-integrating and can deliver up to 38 kilo-basepairs (kb). Adeno-associated viral vectors may integrate or persist as concatemers and have a small physical size with a packaging capacity of 1.6 kb. Herpes simplex viral vectors do not integrate and have a very large capacity of up to 150 kb. Newer adeno-and herpes viral vectors are less immunogenic than earlier generations. Retroviral vectors have a packaging capacity of 7 kb and only transduce dividing cells into which they permanently integrate. Lentiviruses belong to the family of retroviridae but have evolved a nuclear import mechanism for nondividing cells. Derived vectors have a capacity of 7 kb and permanently integrate into both dividing and nondividing cells. All vectors shown here, but not the adeno-associated and retroviral vector, are capable of delivering complex genes that include introns. All of them can deliver exons and small, non-coding RNA (siRNA).

Viruses evolved highly efficient mechanisms of gene transfer and immune evasion a long time ago. The simian immunodeficiency virus, for instance, can be traced back more than 10 million years (Compton et al., 2013) while retroviral orthologues predating the divergence of placental mammals exceed 100 million years (Lee et al., 2013). The downside is that host defense mechanisms are equally mature and can result in pronounced immune reactions in addition to innate restriction at the post-entry level (Kajaste-Rudnitski and Naldini, 2015). A prominent example is the shock syndrome, cytokine release, acute respiratory distress, and multiorgan failure that systemically applied adenoviral vectors can cause (Wilson, 2009). Non-viral delivery using physical or chemical methods to transfer exogenous genes into target cells have a favorable safety profile than viral vectors, but with lower transduction efficiency (Zulliger et al., 2015).

### 2.1. Viral gene delivery

#### 2.1.1. Adenoviral vectors (AdV)

Adenoviruses contain linear, double-stranded DNA. Of species A to F that are subdivided into 50 serotypes (Warnock et al., 2011), B and E are the most frequently used ones for vectors in clinical trials (Ginn et al., 2013). AdV can be produced with relative ease at high titers and high purity (Luo et al., 2007). Gutted AdV are more challenging to generate, however (Kreppel, 2014). They can deliver large transgenes of up to 38 kb that can contain full genes including introns (DelloRusso et al., 2002). The trabecular meshwork (TM) and the inner wall of Schlemm’s canal have been successfully transduced without significantly influencing lens transparency and other tissue architecture nearby (Borrás et al., 1998; Lee et al., 2010). AdV vector preps are unlikely to be affected by pseudotransduction, where contaminating cell fragment proteins from producer cells co-pellet, as may happen with non-banded preps of pseudotyped retroviral vectors (Liu et al., 1996). AdV can transduce both dividing and nondividing tissues. Trabecular meshwork transduction can be achieved without a significant impact on IOP (Borrás et al., 1999).

One shortcoming is that most humans have been exposed to serotypes B and E during their lifetime and have preexisting immunity as a result. Expression in dividing cells is also limited to 5 to 50 days (Borrás et al., 1998; Budenz et al., 1995; Giovingo et al., 2013; Vittitow et al., 2002) because AdV does not integrate. In nondividing cells or in spaces with a relative immune privilege, expression can last for up to a year (Loewen et al., 2004b). AdV is often immunogenic due to coexpressed viral proteins (Borrás et al., 2001) which can be addressed by anti-CD44L treatment (Millar et al., 2008), adjusting the titer (Ethier et al., 2004), or vector gutting (DelloRusso et al., 2002). AdV requires handling at biosafety level (BSL) 2. The nonspecific transduction can be made more specific by transcriptional targeting (Gonzalez et al., 2004) or choosing select sub-serotypes (Ueyama et al., 2014).

#### 2.1.2. Adenovirus-associated vectors (AAV)

AAV, an adenoviral helper-dependent, replication-defective parvovirus, can efficiently infect both dividing and nondividing cells with only a mild immune response and no known pathogenicity (Warnock et al., 2011). AAV-derived vectors have a packaging capacity of approximately 4.6 kb. They are relatively difficult to produce but high titers can be achieved (Grieger et al., 2006). There is an established track record for AAV vectors with inherited eye diseases (Bainbridge et al., 2008) but the transduction efficiency of conventional AAV in TM is very low due to the inability to convert single-stranded DNA into double-stranded DNA, which requires DNA polymerase complexes in host cells (Borrás et al., 2006). Transduction efficiency varies among serotypes of scAAV and host species (Bogner et al., 2015). Expression is slower in the outflow tract than other regions and takes approximately 1 week (Borrás et al., 2015). Gene expression is more persistent with self-complementary vectors and has been reported for at least 3.5 months in rat and 2.3 years in monkey TM (Buie et al., 2010). TM transduction is relatively IOP neutral but mild intraocular inflammation may limit expression (Buie et al., 2010). AAV vectors can be handled at BSL 1 but production typically occurs at BSL 2 due to the use of adenoviral components. Although AAV integrates preferentially into chromosome 19, this is not the case for AAV-derived vectors which typically persist as intracellular head to tail concatemers (Yan et al., 2006, 2005), integrate without specificity, or insert into active genes (Nakai et al., 2003).

#### 2.1.3. HSV vectors

HSV is a 152 kb double-stranded DNA virus (Warnock et al., 2011). HSV-1 based vectors are more commonly used for CNS gene therapy targets than HSV-2 due to the tropism of the HSV-1 wild type (Goins et al., 2014). Wild type HSV-2 is less epileptogenic than HSV-1 yet can also be used for CNS gene therapy and can, for instance, use an intranasal delivery route to treat seizures in a model (Laing et al., 2006). The neurotropism and long latency of HSV may benefit HSV-based vectors, especially for gene transfer to neurons (Chattopadhyay et al., 2005). The packaging capacity of 150 kb is very large (Marconi et al., 2015). First generation HSV vectors can cause severe intraocular inflammation in primates (Liu et al., 1999), but less so in rodents (Spencer et al., 2000). Ninety percent of humans already have antibodies to HSV-1 or HSV-2 (Wald and Corey, 2011). In nonhuman primates, nearly 100% of the TM and non-pigmented ciliary epithelium cells were transduced (Liu et al., 1999), but achieved only 10%-20% transduction efficiency in rats and none in mice following an intracameral bolus (Spencer et al., 2000). While early generation vectors had short term expression measured in days (Liu et al., 1999), long-term expression has been achieved in the CNS with later generation vectors (Zhang et al., 2012). New HSV vectors that are devoid of all five immediate-early genes appear to be able to achieve longer-term expression also in non-neuronal cells (Miyagawa et al., 2015). HSV is handled as BSL 2.

#### 2.1.4. Retroviral vectors (RV)

Retroviruses contain a single-stranded RNA genome of 7 to 12 kb length, which is converted into DNA by reverse transcriptase to linearly integrate into the host genome (Coffin et al., n.d.). The historical classification into type-A, -B, -C and -D is based on their morphology in electron microscopy and is of limited use today. All retroviruses depend on dissolution of the nuclear membrane during cell division to obtain access to the chromatin except lentiviruses, which have evolved an import mechanism. The most commonly used RV is based on a type-C oncovirus, the Moloney murine leukemia virus (MLV), with a simple genome and a packaging capacity of about 7 kb (Nayerossadat et al., 2012). The normal wild type genome length cannot be exceeded in a vector that consists of a minimized viral backbone and a transgene. Expression of transgenes is driven by long terminal repeats (LTRs) that contain splice donors susceptible to cryptic splicing (Armentano et al., 1987). Only dividing cells can be transduced (Lewis and Emerman, 1994) which can be exploited to target proliferative processes (Kimura et al., 1996). MLV appears to favor integration in close proximity to transcription start sites, which may be oncogenic if this occurs near a proto-oncogene or disrupts a tumor suppressor gene (Wu et al., 2003).

#### 2.1.5. Lentiviral vectors (LV)

Lentiviruses belong to the family of retroviruses but have a more complex genome (Warnock et al., 2011) with a packaging capacity of about 7 kb. LV genomically integrate into nondividing cells such as neurons (Loewen et al., 2004b) or nondividing, terminally differentiated cells, that represent the vast majority of cells in an organism, and include the TM (Fig. 2 (Loewen et al., 2001)) or the retinal pigment epithelium (Loewen et al., 2004b, 2003). There is only one clinical trial that uses lentiviral vectors, a phase 1 dose escalation safety study of a subretinally injected vector that expresses endostatin and angiostatin for advanced neovascular age-related macular degeneration (US-1061). Rev restricts RNA splicing in lentiviruses and thereby allows transfer of nucleic acid with cryptic splice sites and complex genes containing introns (Puthenveetil et al., 2004). LVs have been derived from many species and include the human (HIV (Naldini et al., 1996)), simian (SIV (Mangeot et al., 2000)), bovine (BIV (R. Berkowitz et al., 2001)) and feline immunodeficiency virus (FIV (Poeschla et al., 1998)), as well as the equine infectious anemia virus (EIAV (Olsen, 1998)) and visna virus (R. D. Berkowitz et al., 2001). Because the LTRs of HIV vectors are active promoters in human cells, a self-inactivating modification (SIN) was developed (Miyoshi et al., 1998). While 3′-LTR promoter activity could lead to expression of open reading frames downstream of the insertion site, 5′-LTR promoter activity may generate antigenic peptides from Δgag. The U3 deletion of the 3′-LTR SIN design is copied to the 5′-LTR during reverse transcription and inactivates both LTRs of the integrated vector (Loewen and Poeschla, 2005). FIV LTR promoter function is minimal in non-feline cells and any residual promoter activity from either 5′-or 3′-LTR can be avoided by deleting the 172 bp of the U3 Region of the 3′LTR (including the TATA box and binding sites for transcription factors NFκB, NF-ATc and SP1). FIV vectors are typically produced by transient transfection of a three plasmid system (Saenz et al., 2006) while packaging cell lines as well as four plasmid systems are available for HIV vectors (Barde et al., 2010). Insulator elements can be used to prevent silencing (Romero et al., 2015). There may be low-level expression from non-integrating, circular episomal elements that occur after cell entry (Saenz et al., 2004; Yáñez-Muñoz et al., 2006). Non-primate lentiviral vectors are handled at BSL 2 but HIV based vectors require BSL 2+ at most institutions.

**Figure 2.**
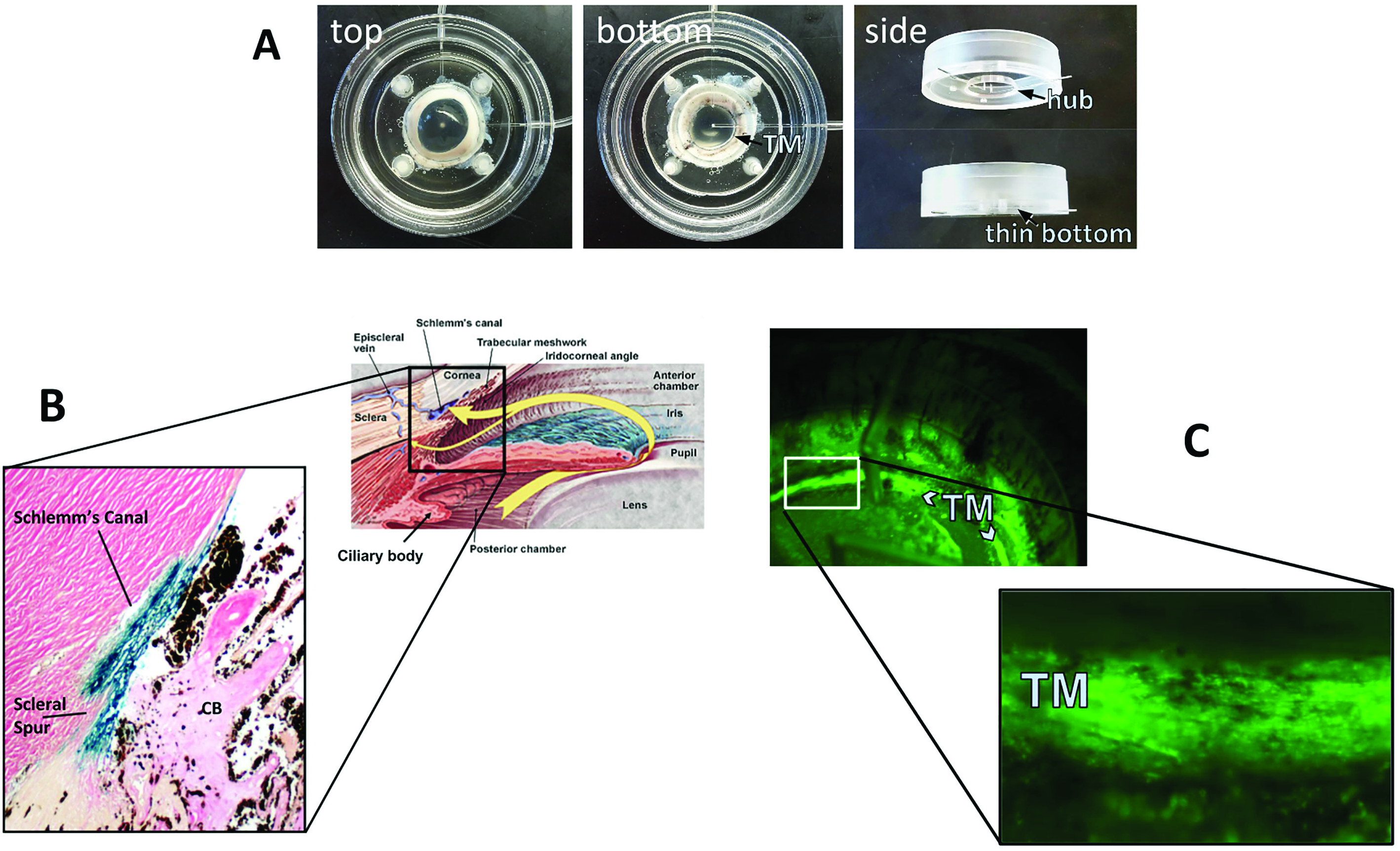
Fig. 2. Anterior segment perfusion culture system (Loewen et al., 2016) (A). FIV transduction of human (B) and porcine eyes (C). A redesigned culture bottom allows direct observation of vector function through the bottom of the dish (C).

FIV and HIV vectors have a high TM transduction efficiency in perfusion-cultured human eyes (Fig. 2 B, (Loewen et al., 2001)) and pig eyes (Fig. 2 C (Loewen et al., 2016)) when compared to transducing units-adjusted Ad vector and MLV with a low cytotoxic effect and immunogenicity (Barraza et al., 2009; Challa et al., 2005; Loewen et al., 2004b). Only transient (Barraza et al., 2009) or no (Barraza et al., 2010; Challa et al., 2005; Zhang et al., 2014) inflammation was recorded. Long-term expression of 455 days in monkeys (Barraza et al., 2009) and 840 days in domestic cats (Khare et al., 2008) has been and serially followed by live gonioscopy (Fig. 3). A temporary decrease of outflow facility was seen in human anterior segment perfusion cultures (Loewen et al., 2002; Zhang et al., 2014) but not in cats (Khare et al., 2008; Loewen et al., 2004a).

**Figure 3.**
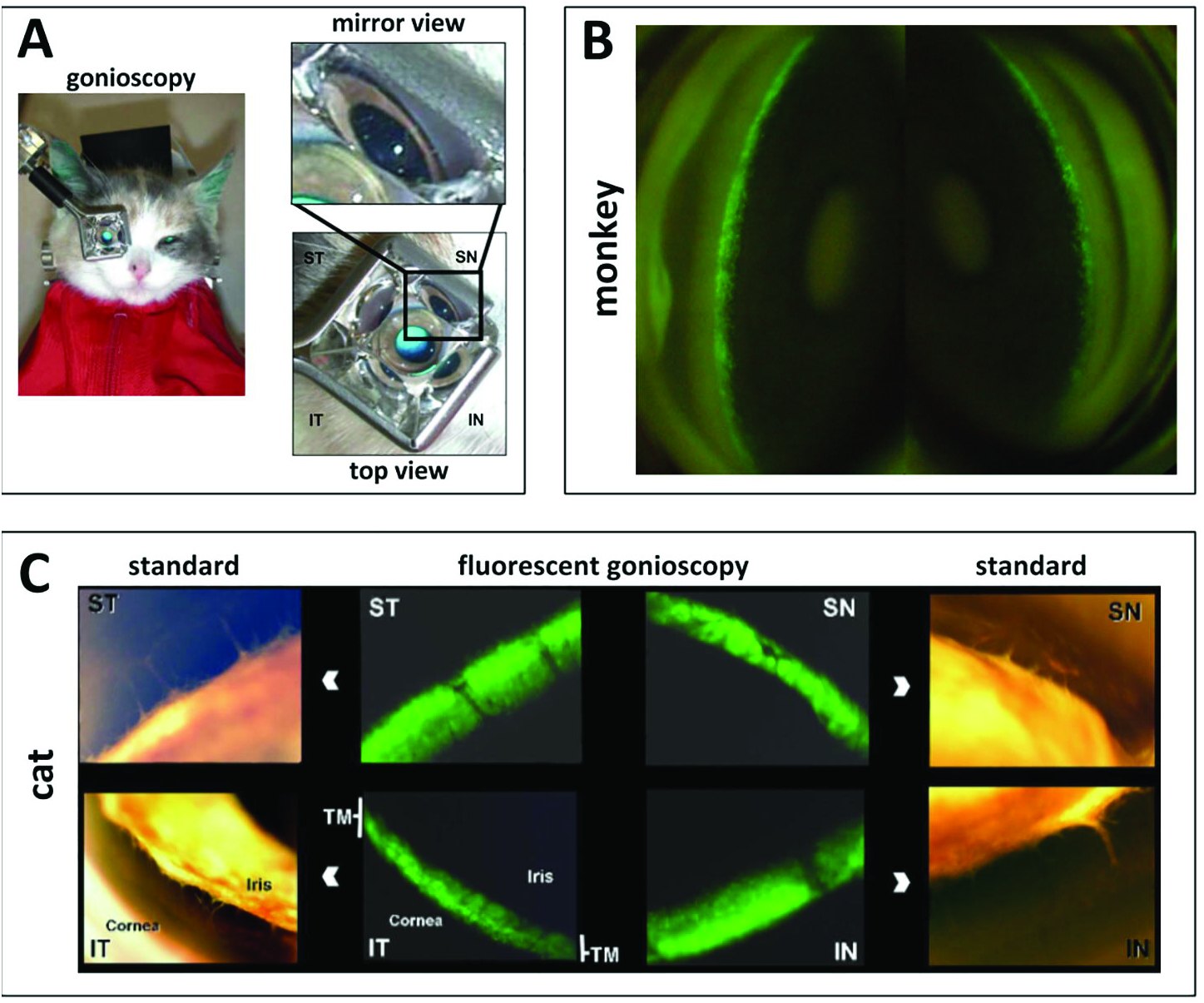
Fig. 3. Live observation of transgene expression with a standard goniolens. Setup for observation of mid-sized animals (A). Serial observation of EGFP marker gene expression in a macaque eye following an intracameral FIV bolus (B (Barraza et al., 2009)). Relatively selective TM transduction and marker gene in the domestic cat (C (Loewen et al., 2004a)).

## 3. Non-viral gene delivery

Nucleic acid can also be delivered by injection or a combination of chemical and physical methods. Advantages are low immunogenicity, low mutagenicity, and large capacity. The limitations are that such transfection is transient and inefficient (Conley et al., 2008).

Naked siRNA can suppress targets in human TM cells. Function is limited to about 48 hours (Comes and Borrás, 2007; Gonzalez and Tan, 2014) and systemic exposure can be detected (Lawrence et al., 2011). Transfection of naked DNA to the outflow tract has not been described.

Nanoparticle can allow non-invasive gene delivery to ocular tissues at greater efficacy than naked DNA (Eljarrat-Binstock et al., 2008). Expression can be detected as early as 4 hours (Losa et al., 1991) and is longer lasting in inner than in outer ocular tissues (Eljarrat-Binstock et al., 2008). Intracameral injection allows transfection of TM (Farjo et al., 2006; Zou et al., 2011). Intravitreal injection of poly (l-lactic acid), polystyrene, and Poly N-isopropylacrylamide particles caused a substantial reduction of IOP and TM inflammation making hyaluronic acid a better candidate for TM gene delivery (Zou et al., 2011).

The sleeping beauty transposon system was first developed 20 years ago (Ivics et al., 1997). It consists of a transposon with the transgene expression cassette and the transposase which can bind to inverted and direct repeats in transposons to facilitate integration. Advantages are long-term expression, high transfection efficiency (Johnen et al., 2012), low pathogenicity, and integration independent of the cell cycle (Ivics and Izsvák, 2011). Electroporation (Johnen et al., 2012), transfection reagents (Belur et al., 2007), and hydrodynamic injection (Bell et al., 2007) do not appear to be effective methods of delivering transposon elements in the anterior segment. To address this problem, a hybrid vector combining a chimeric virus and transposons has been developed (de Silva et al., 2010). EGFP expression was seen in RPE, corneal endothelial, and iris cells after subretinal injection of such hybrid vector in mice (Turunen et al., 2014). However, the advantages provided by hybrid vector should be weighed against a low cost-efficient ratio and the complexity of vector production (Aronovich et al., 2011).

Replicating episomal vectors, which include small circular vectors and artificial chromosomes, can provide sustained gene expression (Zehnpfennig et al., 2010) and avoid unpredictable consequences of gene integration (Lipps et al., 2003). Another possibility is the use of bacteriophage ΦC31 integrase and the attB recognition sequence to enable genomic integration and prolonged expression (at least 4.5 months in mouse retina (Chalberg et al., 2005)). This system has to be electroporated which often causes cataracts, inflammation, and nanophthalmos (Chalberg et al., 2005).

## 4. Applications

### 4.1. Enhancing outflow

Choosing a transgene for gene therapy of glaucoma is difficult because in only less than 10% of glaucoma patients can a single gene be identified to be at fault (Fan et al., 2006): MYOC (Stone et al., 1997), CYP1B1 (Lim et al., 2013), CAV1/CAV2 (Thorleifsson et al., 2010), LTBP2 (Ali et al., 2009), GAS7,TMCO1 (van Koolwijk et al., 2012), and ARHGEF12 (Springelkamp et al., 2015) can all alter outflow. In addition, the conventional outflow tract is a complex structure that consists of many distinctly different components and cell types: the uveoscleral, corneoscleral and deeper juxtacanalicular TM, the inner and outer wall of Schlemm’s canal, collector channels and aqueous veins. These structures have different transduction properties and kinetics: the TM is a phagocytotically active tissue that consists primarily of terminally differentiated, nondividing cells which facilitates transduction with VSV-G pseudotyped lentiviral vectors while the directly adjacent cornea is less transduced when exposed to the same vector bolus (Loewen et al., 2001). The ability of the TM and Schlemm’s canal to present antigens and induce tolerance (anterior chamber associated immune deviation, ACAID, (Stein-Streilein and Streilein, 2002), may work in favor of gene transfer to the outflow tract by reducing immunity against vector and transgene. Schlemm’s canal has both typical and lymphatic vascular properties (Kizhatil et al., 2014) from which collector channels start to sprout during embryogenesis (Ramírez et al., 2004). An application of gene therapeutic substances to the anterior chamber, Schlemm’s canal, or the collector channels has potential to cause a systemic exposure since they drain into the venous circulation.

A well-studied therapeutic target to increase outflow facility is the RhoA and Rho-kinase pathway, which modulates the actin cytoskeleton, cell adhesive interactions, ECM formation, and TM contraction (Wang et al., 2013; Zhang et al., 2008). Increased outflow facility after dominant-negative RhoA or exoenzyme C3 (Rho GTPase inhibitor) by an adenoviral vector was observed in a primate anterior segment perfusion model (Liu et al., 2005; Vittitow et al., 2002). Borras et al used a scAAV-mediated RhoA transfer to prevented elevation of IOP for more than 4 weeks in rats (Borrás et al., 2015). Rho-kinase, a key downstream effector of activated RhoA, may present as a more effective therapeutic target since specific inhibition of this gene increased the outflow facility much greater than inhibiting RhoA (80% vs. 32.5%) (Rao et al., 2005; Vittitow et al., 2002). Barraza et al transduced components (receptor and enzyme) of the prostaglandin pathway using FIV to reduce IOP in domestic cats, an approach that involved codon optimization to avoid rapid RNA degradation of the expressed transgenes (Fig. 4A (Barraza et al., 2010)). Kumar and colleagues found that transduction of tissue plasminogen activator by an adenoviral vector increased outflow facility by 60% in steroid induced glaucoma (Kumar et al., 2013). Herpes simplex thymidine kinase/ganciclovir based TM ablation using conditionally cytotoxic FIV vectors reduced IOP in a rodent model (Fig. 4B (Zhang et al., 2014)). Naked glucocorticoid receptor small interfering RNA (siRNA) in a human anterior segment perfusion model caused significant reduction of MYOC and CDT6 expression by 98% and 85%, respectively (Comes and Borrás, 2007). Similarly, a recent study by Gonzalez and Tan also found that a low dose of siRNA significantly inhibited targeted gene by 53.9% (Gonzalez and Tan, 2014). Silencing ChGn, an enzyme specific to chondroitin sulfate glycosaminoglycan biosynthesis, and caveolin-1 by shRNA caused increased outflow facility in the same model (Aga et al., 2014; Keller et al., 2011). Matrix metalloproteinases play an important role in degrading ECM and decrease the resistance of outflow (De Groef et al., 2013). Gerometta et al (Gerometta et al., 2010) and Spiga et al (Spiga and Borrás, 2010) developed a glucocorticoid-inducible metalloproteinase 1 Ad vector to reduce IOP in the presence of continued corticosteroid application.

**Figure 4.**
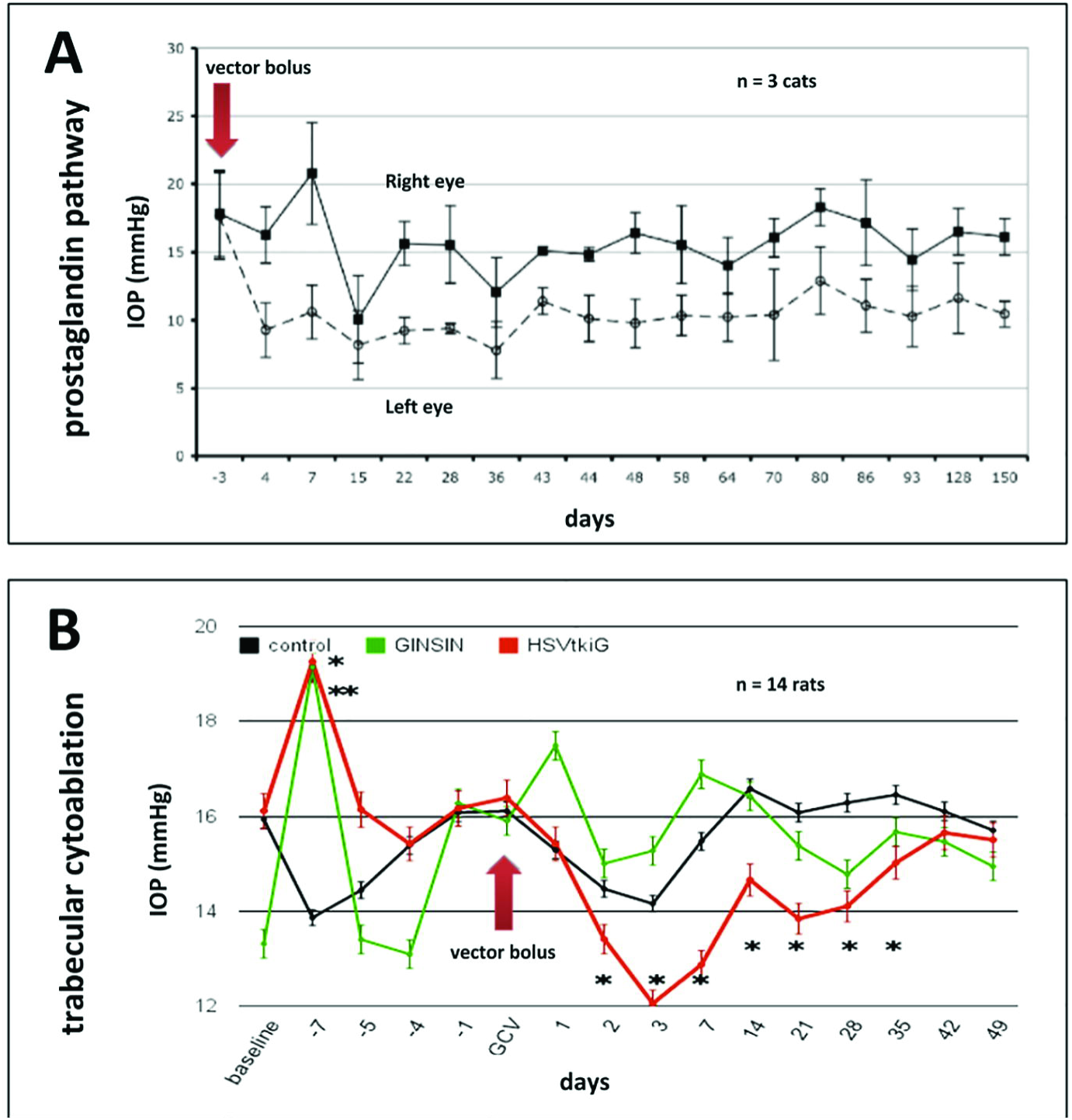
Fig. 4. Examples of gene therapeutic IOP lowering. Prostaglandin pathway transduction shows a long-term IOP reduction after a single vector bolus in the cat (A (Barraza et al., 2010)). Inducible cytoablative vector causes a rapid TM cell loss followed by a recovery of IOP and TM cell numbers (Zhang et al., 2014) in this model of TM regeneration (B).

### 4.2. Generating disease models

Gene transfer to the outflow tract can be used to develop OAG disease models. Members of the TGF-β superfamily (Buie et al., 2013; Robertson et al., 2010) and their downstream effectors (Junglas et al., 2012; Lee et al., 2010; Oh et al., 2013) can induce ECM remodeling in the TM (Fuchshofer and Tamm, 2012; Wordinger et al., 2014) and increase IOP. AdV-mediated overexpression of TGF-β1 and TGF-β2 resulted in reduced outflow facility and ocular hypertension in a dose-dependent manner (Robertson et al., 2010; Shepard et al., 2010). Transduction of BMP2 into TM also increased IOP in rats (Buie et al., 2013) as did transduction of downstream effectors COCH (Lee et al., 2010), CTGF (Junglas et al., 2012), and SPARC (Oh et al., 2013). IOP increased in a dose-dependent fashion following intravitreal injection of adenoviral vector encoding sFRP1 (Wang et al., 2008) and sCD44 (Giovingo et al., 2013) in mice. Lentiviral shRNA silencing was used to investigate the role of three different hyaluronan genes in outflow in human eyes (Keller et al., 2012). Similar methods have identified various roles of versican, a large aggregating CS proteoglycan (Keller et al., 2011), tenascin, a matricellular glycoprotein (Keller et al., 2013), and different caveolins on aqueous outflow resistance and ECM remodeling (Aga et al., 2014).

## 5. Current Challenges and Future Directions

### 5.1. New promoters, models, transgenes

Because cell type specific promoters are not always known, the highly expressing CMV or EF1-α promoters are most commonly used, which has led to expression not only in the TM but also occasionally in corneal endothelial cells (Challa et al., 2005; Zhang et al., 2014) and non-pigmented ciliary epithelial cells (Barraza et al., 2010; Bogner et al., 2015). A TM preferential promoter is the matrix Gla protein promoter (Gonzalez et al., 2004) or chitinase 3-like 1 promoter (Liton et al., 2005). Ueyama et al found that an unmodified Ad35 serotype preferentially transduces TM cells (Ueyama et al., 2014).

To establish a glaucoma animal model by tissue specific transgenesis using viral vectors, other transgenes should be considered. For instance, smad3 is such a candidate that amplifies the TGF-β signal and is essential for TGF-β2 induced ocular hypertension in mice (McDowell et al., 2013). Similarly, Gremlin could be tested as it blocks the negative effect of BMP-4 on TGF-β induced TM fibrosis (Wordinger et al., 2007). S1P_2_ receptor activation has also shown significant increases in the conventional outflow resistance of human and porcine eyes (Sumida and Stamer, 2011) and might allow to create a glaucoma model in those eyes.

Conversely, BMP-4 (Wordinger et al., 2007), Smad7 (Fuchshofer et al., 2009), and BMP7 (Fuchshofer et al., 2007) can antagonize TGF-β mediated TM changes and may be suitable for glaucoma gene therapy. Nitric oxide is an intracellular signaling molecule with IOP-lowering effects (Cavet et al., 2014). Overexpression of endogenous NO synthases (Stamer et al., 2011) and Ghrelin (upstream modulator of NO) (Azevedo-Pinto et al., 2013) could decrease IOP and increase pressure dependent drainage. Transduction of eNOS and Ghrelin into the outflow tract may be a viable strategy for glaucoma therapy. If outflow resistance is located downstream of the TM, overexpression in TM cells may not be sufficient.

### 5.2. New targets and delivery methods

#### 5.2.1. Targets downstream of the trabecular meshwork

The majority of outflow resistance is produced by the extracellular matrix in the juxtacanalicular TM and possibly the inner wall endothelium of Schlemm’s canal while the larger pores of the outer meshwork are not able to generate a significant resistance. In theory, a single pore of 100 micrometer length and a diameter of 20 micrometers may allow for a pressure drop of 5 mmHg (Johnson and Erickson, 2000). It has previously been thought that the aqueous veins would not be able to produce a significant outflow resistance (estimated to be 20 vessels of 50 micrometer diameter and an average length of 1 mm (Rosenquist et al., 1989)). However, this contradicts a mounting body of both experimental (Schuman et al., 1999; Van Buskirk, 1977) and clinical evidence of residual outflow resistance in that after trabecular ablation, IOP remains above 15 mmHg (Bussel et al., 2015, 2014; Kaplowitz et al., 2014). This often overlooked outflow resistance downstream of the TM may provide new treatment opportunities. Although the episcleral venous pressure in normal subjects is between 8 to 10 mmHg (Sultan and Blondeau, 2003), it is estimated to be near 12 mmHg in untreated POAG (Selbach et al., 2005). This matches IOP reduction after trabeculectomy ab interno by 30 to 40% or to 3 to 4 mmHg above that of episcleral venous pressure with an average near 15 to 16 mmHg as evidenced by a formal meta-analysis (Kaplowitz et al., 2016). It is possible that pressure dependent outflow vessel kinking (Francis et al., 2012) and valve-like suspended structures at the collector openings (Martin et al., 2014) have a pressure and flow regulatory function (Hann et al., 2011; Schieber and Toris, 2013) and represent novel structural treatment targets.

As a vessel-like endothelial layer, supplement of endogenous NO in Schlemm’s canal decreased the volume of endothelial cells (Ellis et al., 2010) and increased outflow facility (Chang et al., 2015). A recently identified cytokine signaling pathway may provide additional gene therapy targets that are downstream of the TM (Alvarado et al., 2015). Initiating an expansion of the aqueous vascular bed by reinitiating vessel sprouting may also be feasible. This suggests that transduction of collector channels and Schlemm’s canal might be a possible approach for glaucoma, but only if vectors can make it past the TM. Transduction of the downstream outflow tract is possible using FIV vectors (Fig. 5 C).

**Figure 5.**
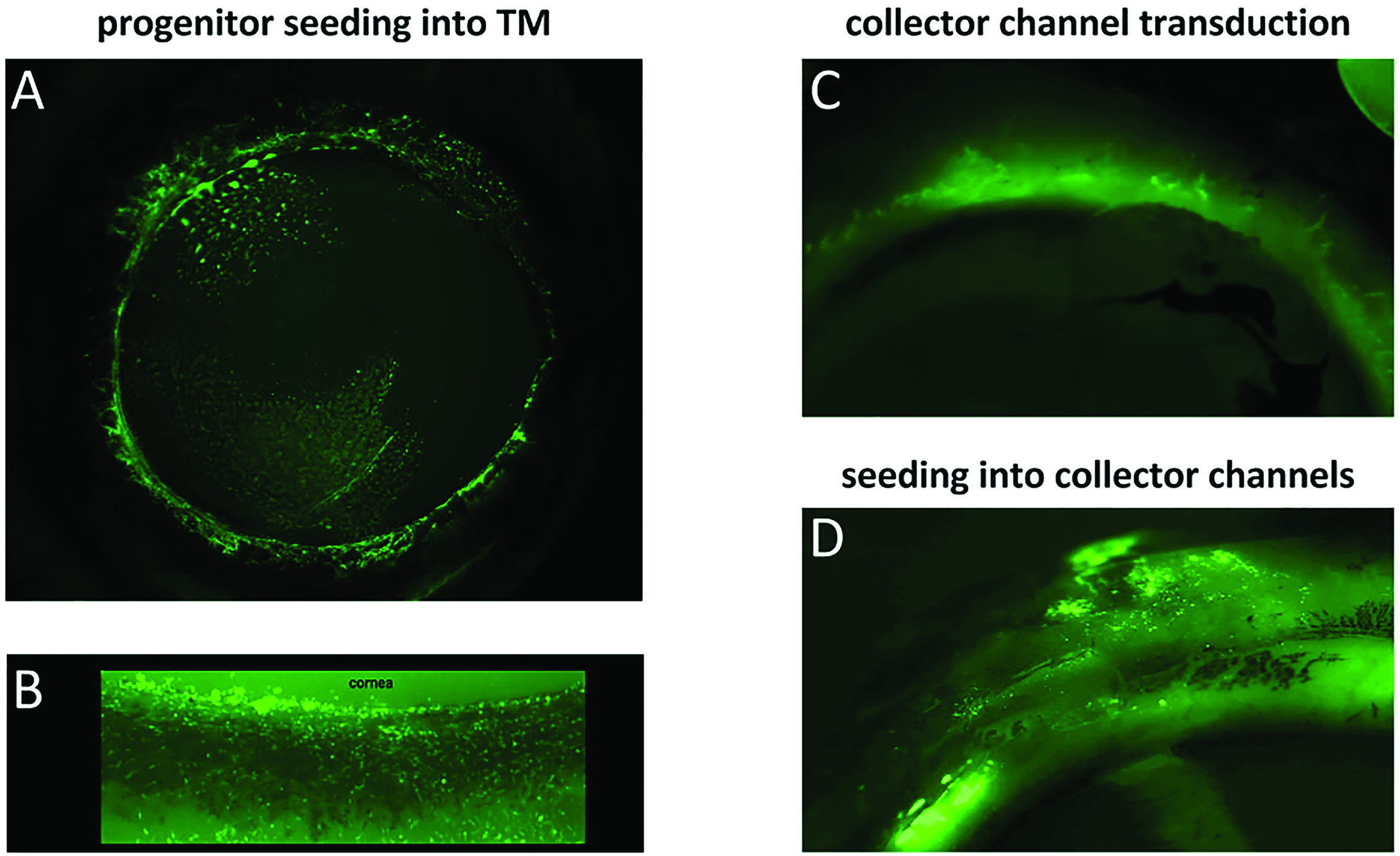
Fig. 5. New cell and gene delivery approaches. (A) Adipose derived tissue stem cells transduced with EGFP expressing FIV vector (Zhang et al., 2014) were seeded into the TM of anterior perfusion cultures of human eyes. View from inside into everted eye. (B) Magnified view of TM from experiment (A). (C) Collector channel transduction with EGFP expressing FIV vector. External, frontal view of perfused porcine eyes (Loewen et al., 2016). An ab interno trabeculectomy (Kaplowitz et al., 2014) was performed followed by an intracameral vector bolus. (D) Cell seeding model with EGFP expressing fibroblasts (Oatts et al., 2013). External, frontal view of perfused porcine eyes. An ab interno trabeculectomy (Kaplowitz et al., 2014) was performed followed by an intracameral cell bolus.

#### 5.2.2. Treatment with engineered cells

As an alternative approach, progenitor or other suitable cells could be engineered in vitro and then seeded into the outflow tract (Fig. 5 A and B) as done in other compartments of the eye (Flachsbarth et al., 2014). Human TM stem cells can be seeded into mouse TM (Du et al., 2013). Cells that are genetically engineered to overexpress therapeutic genes could similarly be transplanted into the TM or into the outflow tract (Fig. 5 D).

#### 5.2.2. Delivery of non-coding RNA

In addition to the long established ribosomal and transfer RNA (tRNA), and siRNA discussed above, several other non-coding RNA (ncRNA) species exist, including micro RNA (miRNA), small nuclear RNA (sn), P-element induced wimpy testis interacting RNA (piwi- or piRNA), as well as long non-coding RNAs (lncRNAs). The major difference between siRNAs and miRNAs is that miRNAs have multiple targets whereas siRNAs are highly specific. MiRNAs can inhibit the contraction of TM cells and ECM short term and up to 10 miRNAs can be delivered by one viral vector (Gonzalez et al., 2014). LncRNAs exceed 200 nucleotides in length, and regulate expression at the transcriptional and post-transcriptional level and make up the majority (about 98%) of the transcriptome (Mercer et al., 2009). LncRNAs may be transcribed as partial or whole natural antisense transcripts (NAT) of coding genes, or they might be located between genes or within introns. Some lncRNAs originate from so-called pseudogenes (Milligan and Lipovich, 2014). They can be described as intergenic, intronic, overlapping, antisense, bidirectional or processed (Mattick and Rinn, 2015; Peschansky and Wahlestedt, 2014). Other characteristics are tissue specific expression, formation of secondary structures, post-transcriptional processing, and mostly nuclear location.

LncRNA MALAT1, implicated in microvascular dysfunction, was recently shown to regulate neurodegeneration and may be involved in glaucoma (Yao et al., 2016). Another study of lncRNA in the eye demonstrated that a myocardial infarction associated transcript (MIAT) knockdown could suppress tumor necrosis factor-alpha-induced abnormal proliferation of human lens epithelial cells (Shen et al., 2016). Promoter activity of a lncRNA encoded on the opposite strand of LOXL1 is influenced by cellular stressors associated with exfoliation syndrome (Hauser et al., 2015). As these examples highlight, lncRNAs may carry out both gene inhibition and gene activation through a range of diverse mechanisms that are made even more complex by the fact that 40% of coding genes have an overlapping antisense transcription. Could it be possible that a highly variable post transcriptional regulation contributes to the range of glaucoma phenotypes? LncRNAs might play a key role because they are poorly conserved between species, and with about 30,000 different transcripts, are estimated to be present in humans at 100 fold the number of messenger RNA. Our view of the human genome and understanding of human diseases is constantly evolving. The research tools and strategies presented here will help to explore cause and effect in the highly complex pathogenesis of glaucoma that has the simple endpoint of elevated IOPs.

## Acknowledgements

Many scientists have contributed to the development of tools and techniques that allow gene transfer and transfer of non-coding material to the outflow tract. Space constraints limited a more complete discussion. Omission of credit was not intended. This work was supported in part the National Institutes of Health research grant EY022737, the Eye and Ear Foundation of Pittsburgh and by the Louis J. Fox Center for Vision Regeneration.

